# Development and validation of a multidimensional composite pain scale for Rabbit (CANCRS) in a clinical environment

**DOI:** 10.1101/728543

**Authors:** Penelope Banchi, Giuseppe Quaranta, Alessandro Ricci, Mitzy Mauthe von Degerfeld

**Affiliations:** C.A.N.C. (Centro Animali Non Convenzionali), Dip. di Scienze Veterinarie, Università degli Studi di Torino, 10095, Grugliasco (TO), Italy; Dip. di Scienze Veterinarie, Università degli Studi di Torino, 10095, Grugliasco (TO), Italy

## Abstract

The main objective of this study was to develop a multidimensional composite pain scale for assessing and quantifying pain in rabbits (CANCRS); to this purpose, Rabbit Grimace Scale (RbtGS) and a scale including clinical parameters (CPS) were merged; the two scales performances were also evaluated individually, in order to validate RbtGS in a clinical setting and to verify clinical parameters usefulness in detecting pain.

Rabbits (n=116) were evaluated by two raters, who could be veterinarians (V) or veterinary medicine students (S). Raters were asked to report the time needed for any evaluation and a total score, in order to define a pain class. Pain classes (No pain, Discomfort, Moderate pain and Severe pain) matched presumptive pain classes and accordingly, the validity of the three scales was measured using Chi-square test.

Patients (n=69) were evaluated by one V and one S, allowing to assess the impact that the experience has on the results. Inter-rater reliability was tested for each scale using intra-class correlation coefficient (ICC) and for each parameter of the CANCRS using Cohen’s kappa. Validity results show that only CANCRS and RbtGS efficiently reveal pain, but both tend to underestimate it.

Inter-rater reliability was very good for both CANCRS and CPS, suggesting that experience has little influence on the results; rater’s experience has a greater impact using RbtGS.

Inter-rater agreement was at least good for each parameter.

In conclusion, CPS alone is neither effective or reliable in quantifying pain.

The RbtGS is a useful tool in clinical practice, when coping with many rabbit breeds; however, training is beneficial for a better use of the scale.

The CANCRS is easy and fast to use and enrich RbtGS with some clinical parameters that should be monitored during any clinical examination, leading to a more exhaustive evaluation of the patient.

## Introduction

Pain has been defined by the International Association for the Study of Pain (IASP) as ‘an unpleasant sensory and emotional experience associated with actual or potential tissue damage’ ^1^. Appropriate pain recognition and treatment represent an ethical obligation for veterinarians in order to preserve the patient’s health and quality of life ^2^.

However, assessment and quantification of pain are arduous, due to great variability of pain expressions among individuals and species. In rabbits pain recognition can be particularly challenging because, as many other prey species, these animals are predisposed to mask any sign of pain^3^. Physiological parameters have limited function as they can be altered in any stressful situation. An increase in heart and respiratory rate can be seen whenever the patient is handled^4,5^, as it happens during clinical examinations. In this context, behavioral changes are not effective for pain assessing and quantification, since a prolonged monitoring is required and there is no evidence that changes observed are related to pain severity^6^. Furthermore, rabbits seem to respond to pain and distress by remaining motionless^6–8^, especially in the presence of an observer.

Multidimensional composite pain scales are available for many species, such as canine and feline^9–12^ and represent a useful tool to conduct an immediate and structured pain quantification. However, there is no validated equivalent for rabbits and the only existing pain scale for this species is the Rabbit Grimace Scale (RbtGS)^13^, which uses changes in facial expression to quantify pain. Grimace scales have been developed and validated in several species, such as Horse^14^, Sheep^15^, Ferrets^16^ and laboratory small rodents^17,18^.

The Rabbit Grimace Scale is based on five facial action units (FAU). Even though its use is faster than assessing several behavioral indicators, the main limitation of RbtGS is that it is based only on New Zealand White (NZW) rabbits and tested in standardized conditions. Furthermore, classes for pain grading and quantification have never been defined, therefore, the scale is not useful to determine whether patients need an analgesic treatment. For these reasons the RbtGS has not yet proven to be suitable for assessing pain in clinical settings^3^.

The objectives of the current study were to develop a multidimensional composite pain scale for the assessment and quantification of pain in rabbits (CANCRS) and to validate the RbtGS in a clinical setting on different rabbit breeds. The former combines the RbtGS with some clinical parameters (CPS). Furthermore, the aim was to define pain classes to achieve improvements in the identification of pain and the need of an analgesic treatment plan.

## Materials and methods

### Animals

One hundred sixteen client-owned rabbits admitted to the C.A.N.C. (the exotics and wild animals Veterinary Teaching Hospital of the Veterinary Science DPT. of University of Torino), were considered for inclusion between the years 2016 and 2018. No restrictions were placed on the breed, sex, age or weight of the rabbits.

Critically ill patients with respiratory distress syndrome or in need of oxygen administration were excluded from the study in order to avoid further stress to the animals. In some patients, facial expression was not evaluable, therefore they were excluded from the study as well. Stuporous and comatose states also represented exclusion criteria, considering stupor as a state of lethargy and immobility with diminished responsiveness to stimulation and coma as a deep state of prolonged unconsciousness and unresponsiveness to external stimuli^19^.

Proper informed consent was collected prior to each clinical evaluation.

### Pain scales

A multidimensional composite pain scale (CANCRS) was developed for rabbits, merging the RbtGS with a Clinical Parameters Scale (CPS), which includes some physiologic data (pupil dilation, respiratory rate, respiratory pattern, heart rate) and behavioral responses (response to palpation, mental status and vocalization).

The RbtGS considers five facial action units (FAU): orbital tightening, cheek flattering, nostril shape, whisker position, ear position. Each FAU was scored according to whether it was not present (score 0), moderately present (score 1) and obviously present (score 2). Lop rabbits ear morphology does not permit to evaluate ear position properly, therefore this parameter was not considered when assessing pain in such breeds.

For what concerns the CPS, the scores were given as follow. In response to noxious stimulation, pupillary dilation reflex occurs^20^; therefore, pupil dilation was included in our composite pain scale and it was scored as present (score 1) or not present (score 0).

Since heart rate is extremely variable and difficult to assess in rabbits^5^, it was asked the raters to score percentage increases of the maximum physiological value (250 bpm)^21^. Increases greater than 20% (score 1) and 50% (score 2) were considered, for increases smaller than 20% the score assigned was 0.

The average respiratory rate for an healthy subject was considered in a values range from 30 to 60 bpm^21^; the possible scores assigned were four: rates of 60 bpm or less (score 0), rates between 61 and 72 (score 1), rates between 73 and 90 (score 2) and rates higher than 90 bpm (score 3). Respiratory pattern was considered as eupneic (score 0) or dyspneic (score 1).

When the suspected painful area was palpated the reaction was recorded and classified as no reaction (score 0), reaction during the palpation (score 1) and reaction before the palpation (score 2).

For what concerns vocalizations, the possibilities were absence of vocalization (score 0), vocalization when touched (score 1), intermittent vocalization without any contact with the operator (score 2) or continuous vocalization (score 3).

The mental status was classified as ‘normal’ (score 0), ‘depression’ (score 1) and ‘obtundation’ (score 2).

The medical history of the patient was reported on each paper in order to establish a presumptive pain (PP) class according to literature^22^, since, as it happens for human beings, it is plausible that pain is directly proportional to the damage’s extension. No pain (NP), discomfort (D), moderate pain (MP) and severe pain (SP) were the four pain classes that were created. Raters were asked to report the time they spent on the evaluation of each rabbit on the printed copy of the scale, in order to establish the average time needed to use the assessment tool. Parameters of the three scales are summarized in *Table 1*.

**Table 1.**
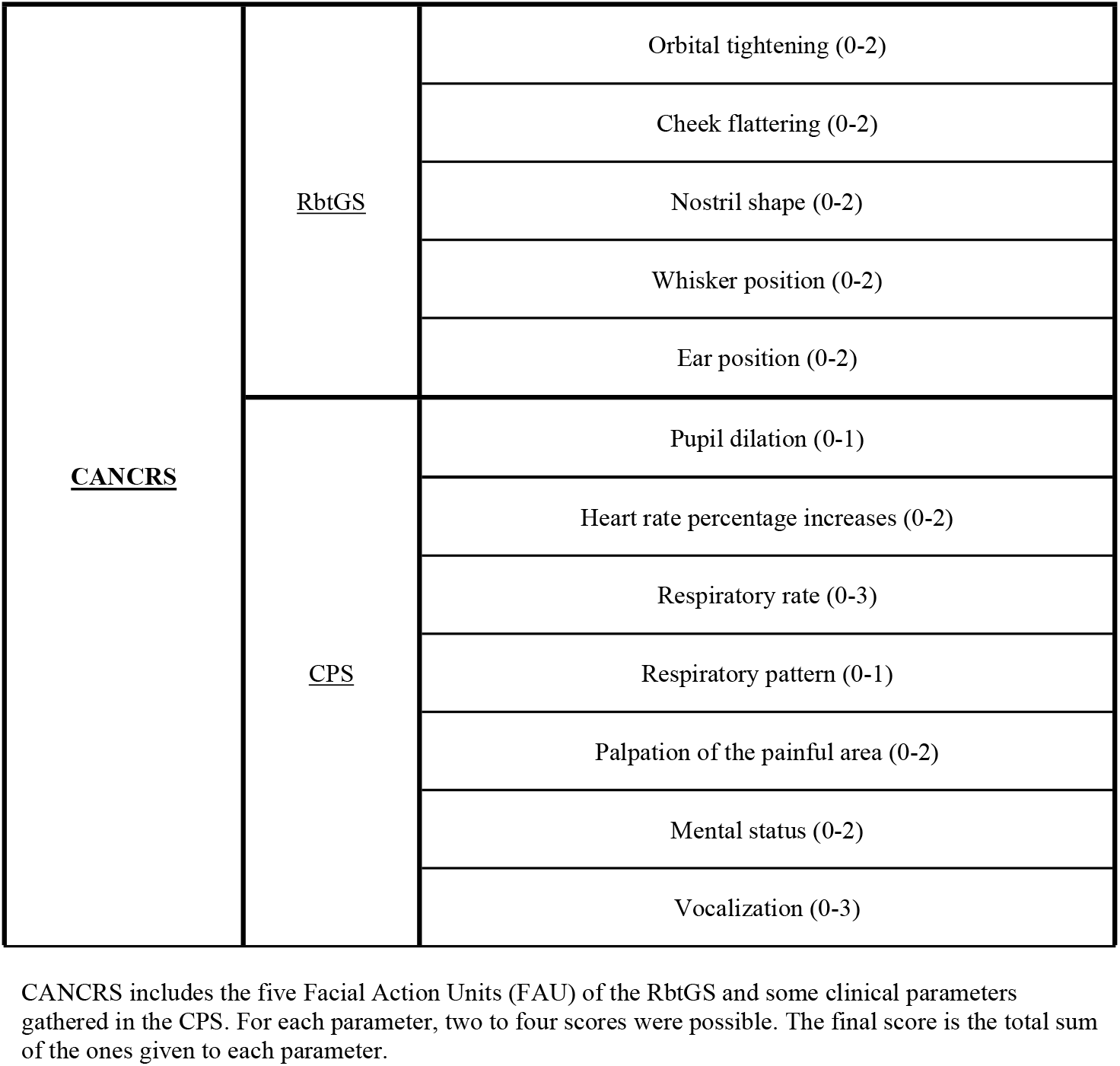
Parameters of the three scales

### Scoring method

Each one of the 116 rabbits included in the study was evaluated independently by two raters during the veterinary clinical examination, allowing to assess the inter-rater reliability; this justify the existence of two documented evaluations (A and B) for each patient. The raters were both veterinarians (V) or a couple made up of a veterinarian and a veterinary medicine students (S), as it happened in sixty-nine cases (56 rabbits and 13 Lop rabbits); this allowed to test the impact that the experience has on the result of the evaluation.

Raters were familiar with using the scale and provided with a printed copy for each rabbit. To ensure consistency, each rater also received clear instructions on how to use the scale and they were supported by the presence of images explaining how to evaluate the FAU considered by the RbtGS. Raters were instructed to first fill in the RbtGS part, assessing the rabbit visually from a distance, prior to opening the cage, interacting with the patient and fill in the second part (CPS) of the scale, which included physiologic and behavioral data.

Each patient was assigned a score for the RbtGS, the CPS and, consequently, for the composite pain scale. Pain scores ranged from 0 to 24 for the CANCRS, from 0 to 14 for the CPS and from 0 to 10 for the RbtGS or from 0 to 8 for the Lop rabbit version of the RbtGS.

### Statistical analyses

The mean of the times required for using the CANCRS was calculated. For each scale, the scores were distributed in the same four classes created for the PP (presumptive pain diagnosis). This allowed the interpretation of the scores as ordinal rather than continuous and to compare the scales with each other and with presumptive pain categories.

### Inter-rater reliability

Sixty-nine rabbits were independently evaluated by a couple made of V and S and were considered to assess inter-rater reliability; to this purpose, intra-class correlation coefficient (ICC) involving a two-way random effect model with 95% confidence intervals was calculated. Furthermore, Cohen’s kappa was determined for each parameter separately, this permitted an estimation of the inter-rater agreement; ear position was assessed in a cohort of 56 rabbits, since 13 patients were Lop rabbits and ear position assessment is not possible on this breed. The results of inter-rater reliability were interpreted using Altman’s classification^23^, in which values ranging from 0.81-1.0, 0.61-0.80, 0.41-0.60, 0.20-0.40 and <0.20 are considered very good, good, moderate, fair, and poor, respectively. Consistently with concurrent recommendations^24^, it was determined a priori that correlations ranging from 0.4-0.8 would be acceptable.

### Validity

Pain assessment was performed on each one of the 116 rabbits included in the study by two raters; the two assessments were randomly allocated into group A or group B; the two groups were considered independently.

The Chi squared test was used to verify how frequently the score assigned by using each scale (CANCRS, CPS and RbtGS) had a correspondence with PP class.

P value < 0.05 was considered significant. Statistical analyses were performed with the software *R* version 3.2.2.

## Results

The mean time required to use the CANCR scale is 209 sec (min 132s-max 341s).

### Inter-rater reliability

The ICC was 0.88 (p < 0.001; 95% CI 0.81–0.92) between V and S for the CANCRS indicating very good inter-rater reliability between veterinarians and students (*Figure 1*).

**Fig. 1.**
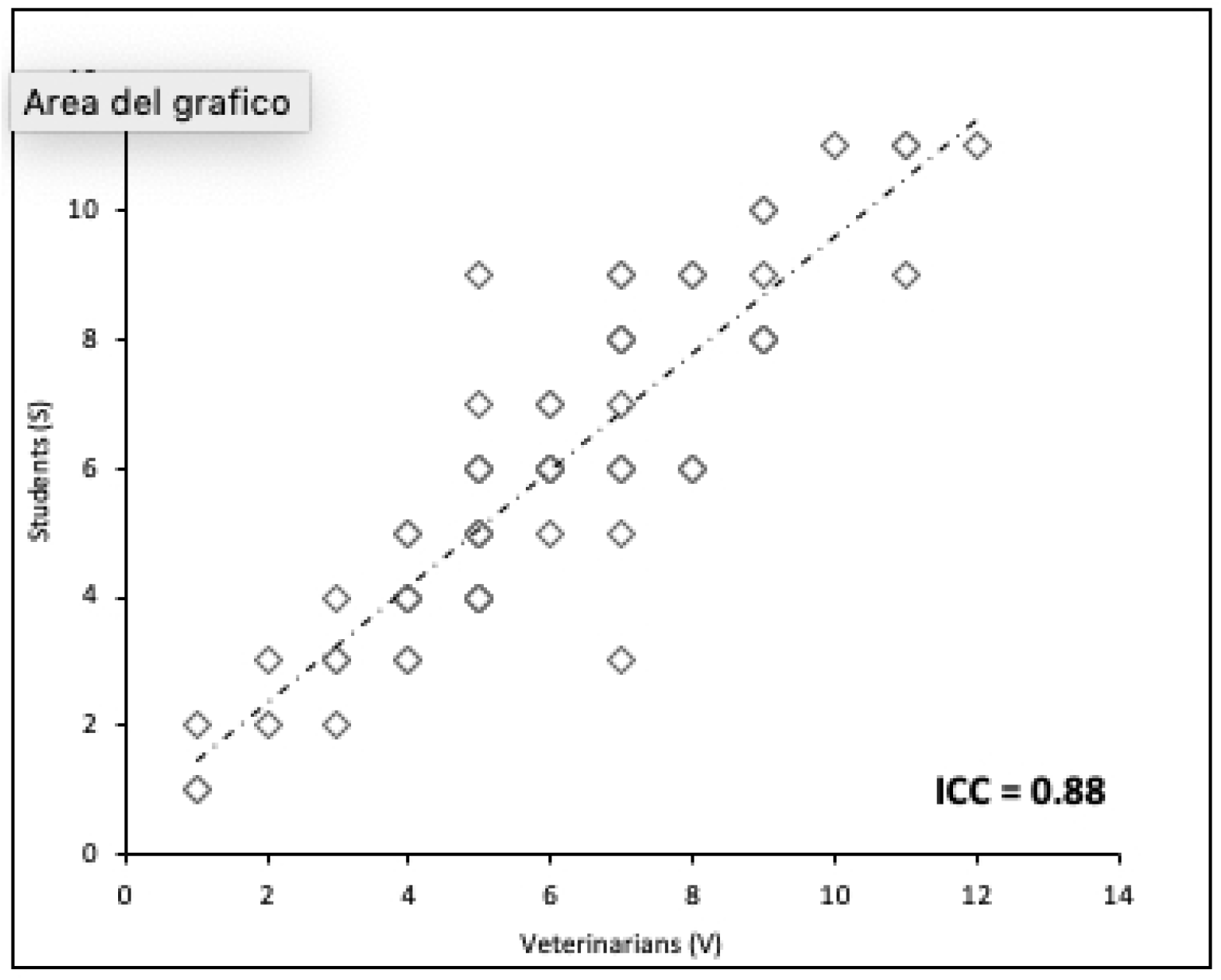
Determination of the inter-rater reliability for the CANCRS. All scores expressed by Veterinary Medicine Students (S) vs scores expressed by Veterinarians (V) are shown in the plot. Unanimous scores are disposed on the dashed trend line. Analysis of the inter-rater reliability revealed an ICC of 0.88, which is considered to be very good, according to Altman’s classification.

The ICC was 0,97 (p < 0.001; 95% CI 0.96–0.98) between V and S for the CPS, indicating very good inter-rater reliability (*Figure 2*).

**Fig. 2.**
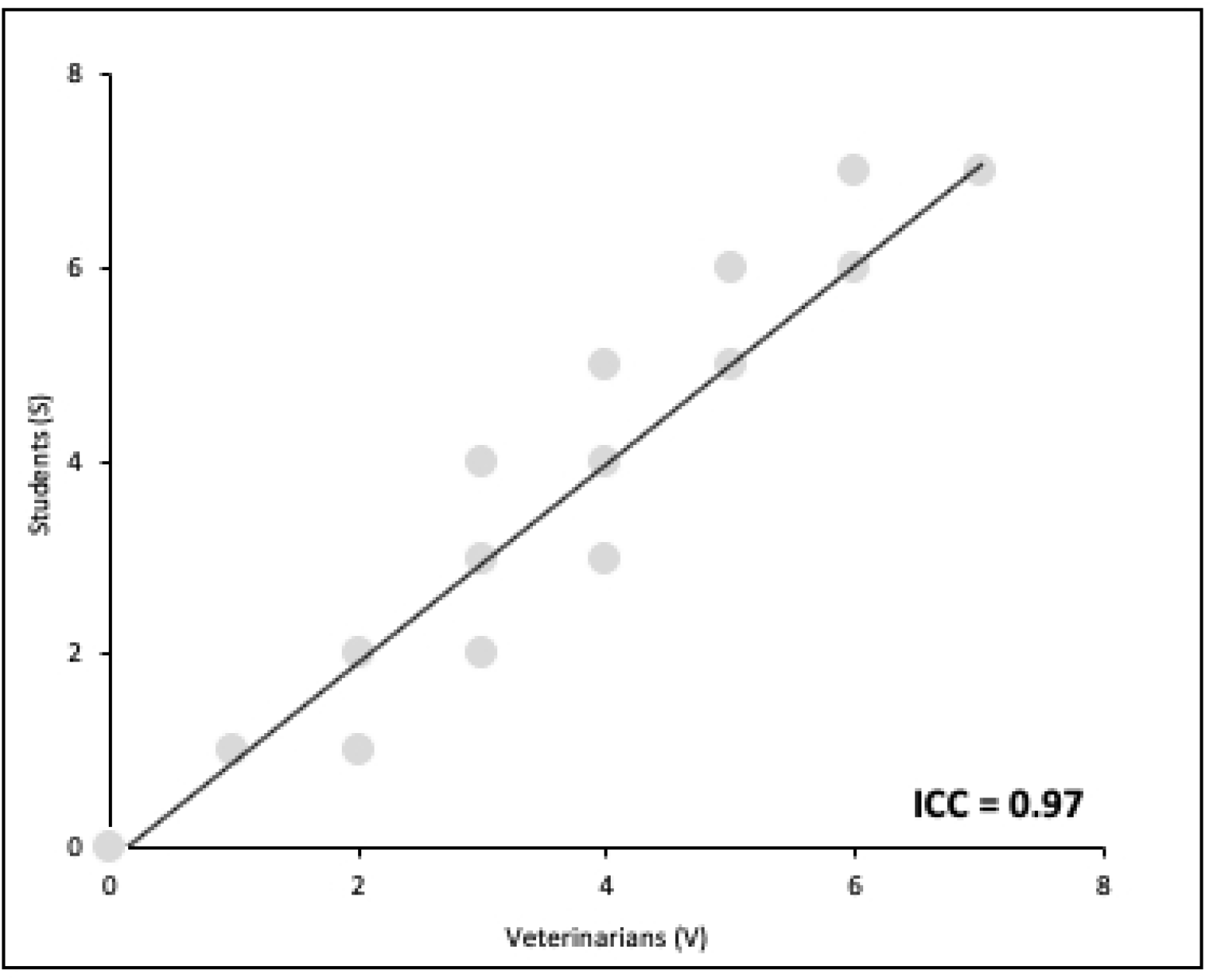
Determination of the inter-rater reliability for the CPS. All scores expressed by Veterinary Medicine Students (S) vs scores expressed by Veterinarians (V) are shown in the plot. Unanimous scores are disposed on the trend line. Analysis of the inter-rater reliability revealed an ICC of 0.97, which is considered to be very good, according to Altman’s classification.

Finally, the ICC was 0.77 (p < 0.001; 95% CI 0.66–0.85) between V and S for the RbtGS, indicating good inter-rater reliability (*Figure 3*)

**Fig. 3.**
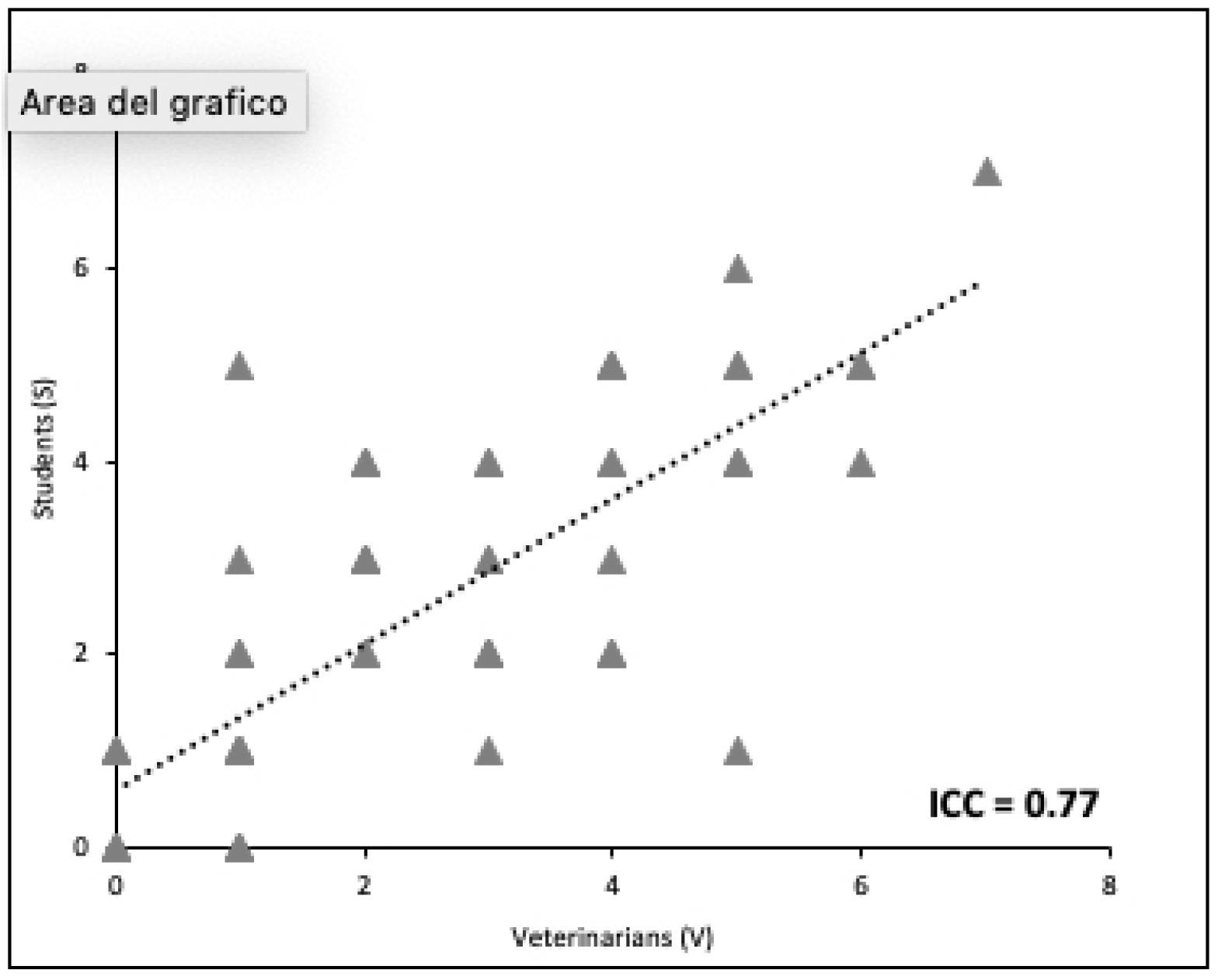
Determination of the inter-rater reliability for the RbtGS. All scores expressed by Veterinary Medicine Students vs scores expressed by Veterinarians are shown in the plot. Unanimous scores are disposed on the dashed trend line. Analysis of the inter-rater reliability revealed an ICC of 0.77, which is considered to be good, according to Altman’s classification.

For what concerns Cohen’s kappa values, acceptable results were obtained for orbital tightening (0.60), cheek flattering (0.47), nostril shape (0.51) and whisker position (0.56). Ear position (0.82), pupil dilation (0.87), respiratory rate (0.96), respiratory pattern (0.89), heart rate (0.94), response to palpation (0.86), mental status (0.91) and vocalization (0.88) resulted to have very good inter-rater agreement.

### Validity

The Chi squared test indicated that the CANCRS-scale and the RbtGS scores are significantly related to PP (P < 0.05), for both group A and B. Otherwise, the test indicated that the CPS score frequencies could be randomly obtained (P > 0.05) for both of the groups (*Figures 4-9*).

**Fig. 4.**
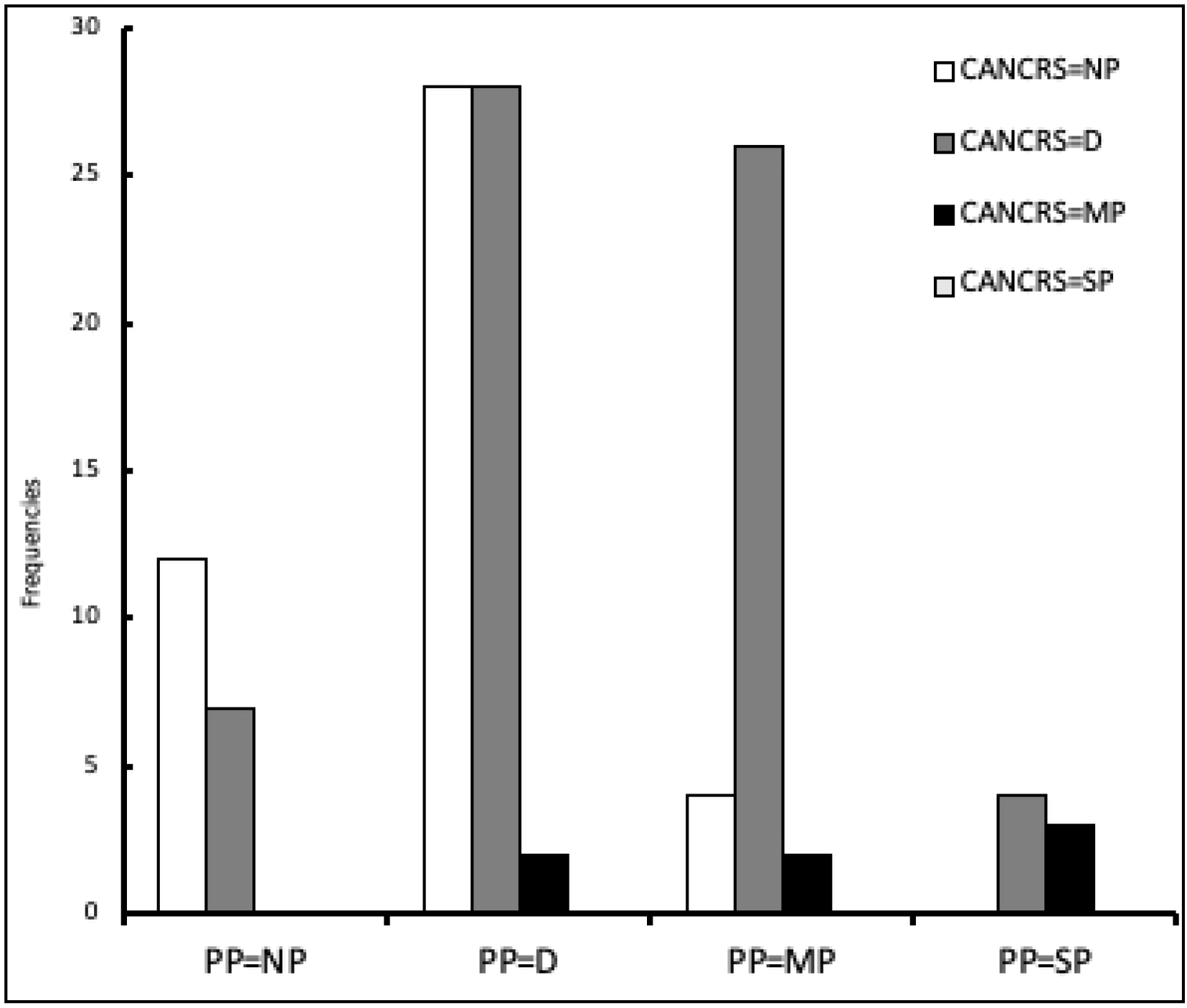
Distribution of the scores resulting from the use of the CANCRS in presumptive pain classes (PP) for group. **A.** Patients in a presumptive condition of pain absence (PP=NP) are mostly classified as NP or D; rabbits in discomfort (PP=D) are mostly classified as NP or D; presumed MP cases are mostly classified as D; none of the presumed cases of several pain was detected. Although, X^2^ test results (p<0.005) show that frequencies are not randomly obtained, but diagnosis obtained by assessing pain with the CANCRS are related to PP.

**Fig. 5.**
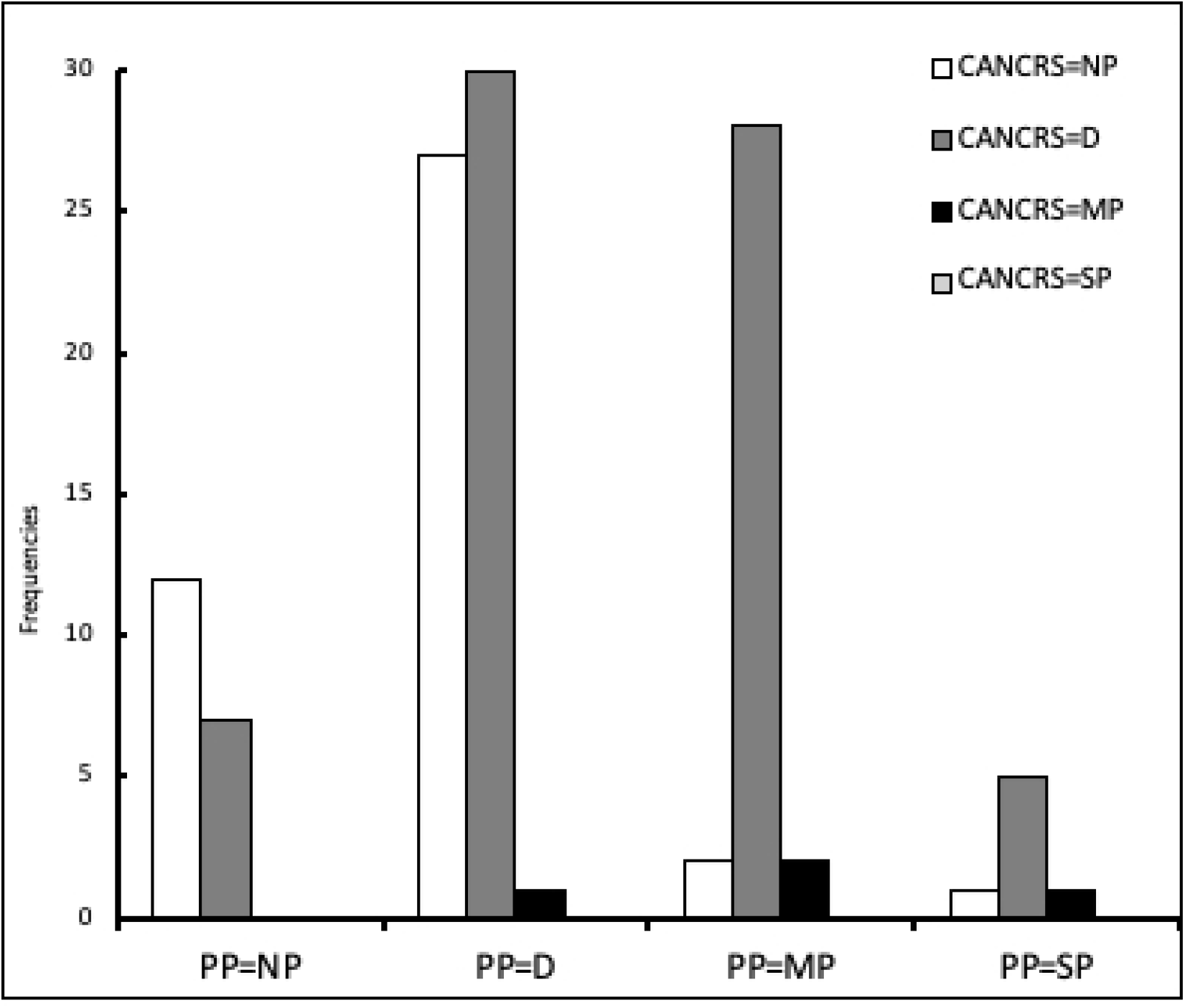
Distribution of the scores resulting from the use of the CANCRS in presumptive pain classes (PP) for group. **B.** Patients in a presumptive condition of pain absence (PP=NP) are mostly classified as NP or D; rabbits in discomfort (PP=D) are mostly classified as D or NP; presumed MP cases are mostly classified as D; none of the presumed cases of several pain was detected. Although, X^2^ test results (p<0.005) show that frequencies are not randomly obtained, but diagnosis obtained by assessing pain with the CANCRS are related to PP.

**Fig. 6.**
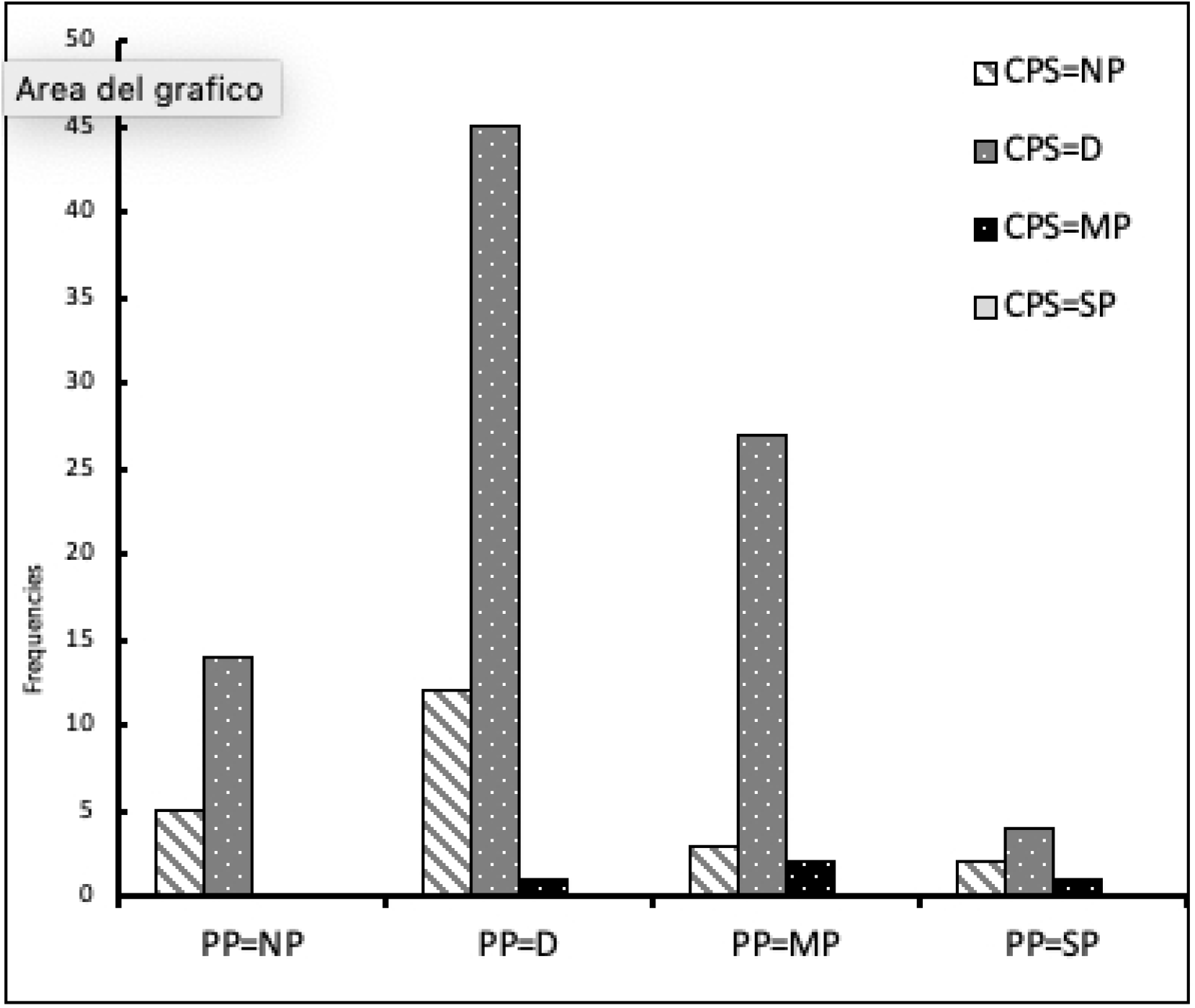
Distribution of the scores resulting from the use of the CPS in presumptive pain classes (PP) for group A. Patients in a presumptive condition of pain absence (PP=NP) are mostly classified as D or NP; rabbits in discomfort (PP=D) are mostly classified as D; presumed MP cases are mostly classified as D; none of the presumed cases of several pain was detected. X^2^ test results (p>0.005) show that frequencies could be randomly obtained, and that there is no relation between CPS and PP.

**Fig. 7.**
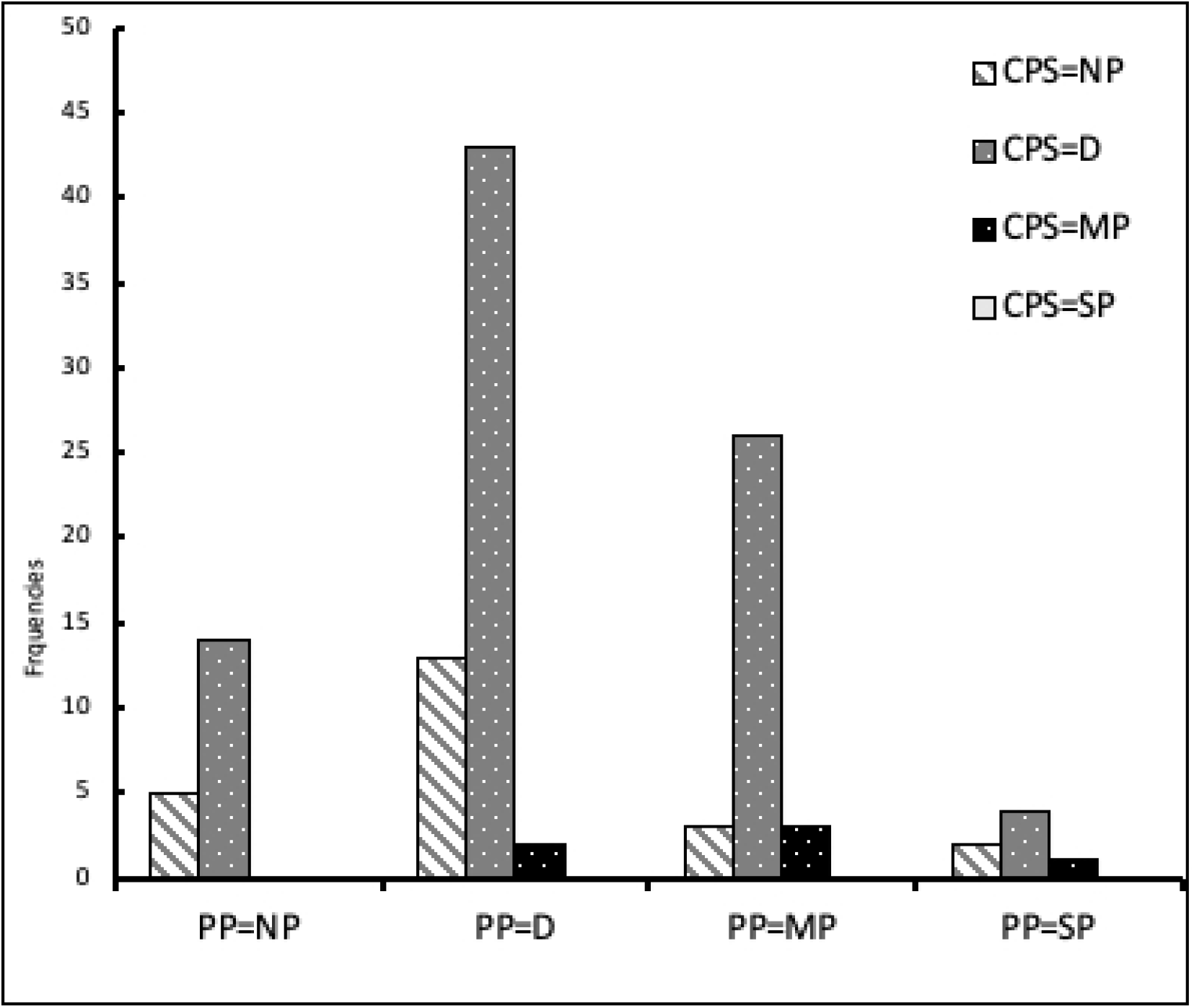
Distribution of the scores resulting from the use of the CPS in presumptive pain classes (PP) for group B. Patients in a presumptive condition of pain absence (PP=NP) are mostly classified as D or NP; rabbits in discomfort (PP=D) are mostly classified as D; presumed MP cases are mostly classified as D; none of the presumed cases of several pain was detected. X^2^ test results (p>0.005) show that frequencies could be randomly obtained, and that there is no relation between CPS and PP.

**Fig. 8.**
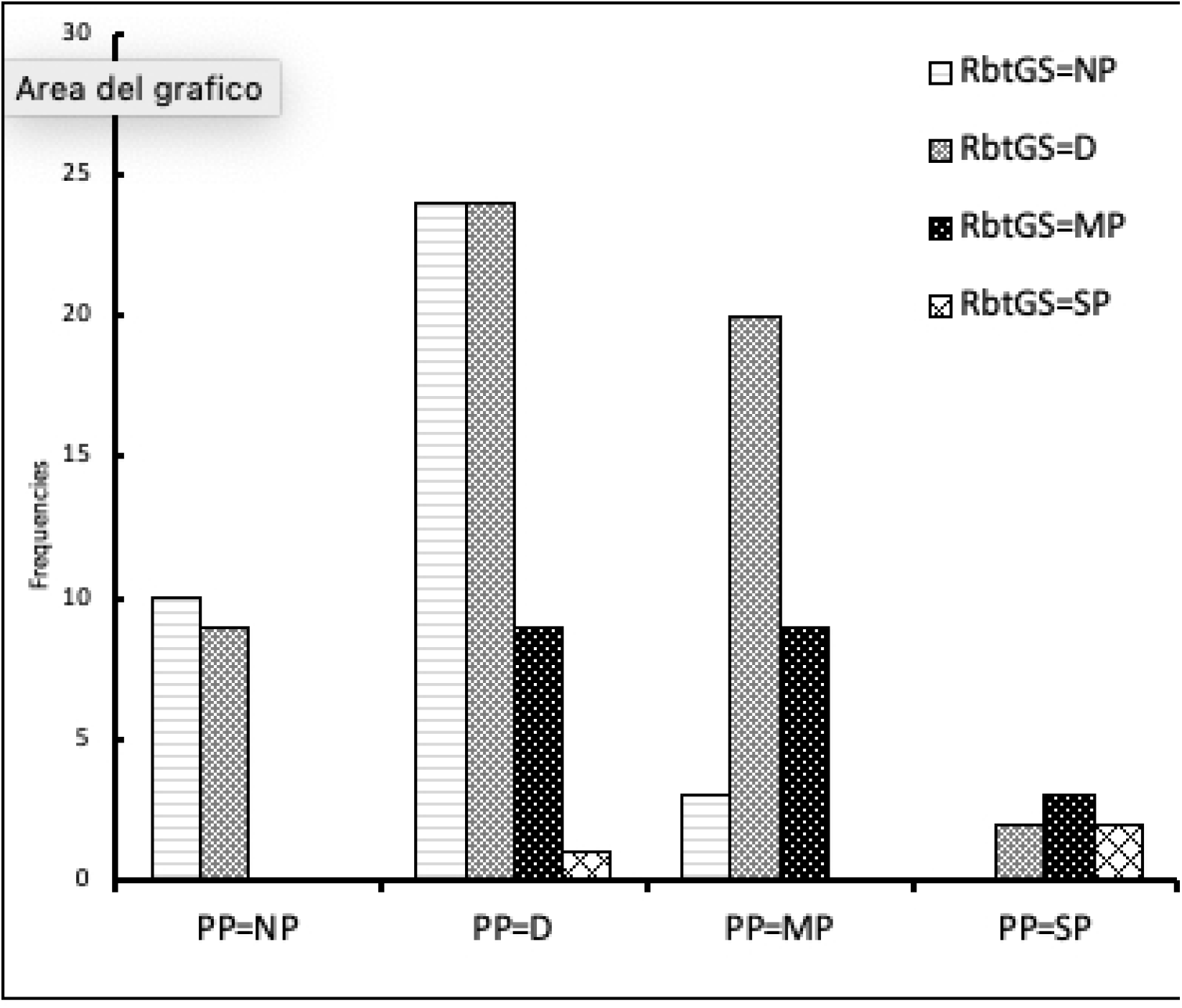
Distribution of the scores resulting from the use of the RbtGS in presumptive pain classes (PP) for group. **A.** Patients in a presumptive condition of pain absence (PP=NP) are mostly classified as NP or D; rabbits in discomfort (PP=D) are equally classified as NP or D; presumed MP cases are mostly classified as D; the RbtGS was the only tool that was able to detect some cases of presumed several pain. X^2^ test results (p<0.005) show that frequencies are not randomly obtained, but diagnosis obtained by assessing pain with the RbtGS are related to PP.

**Fig. 9.**
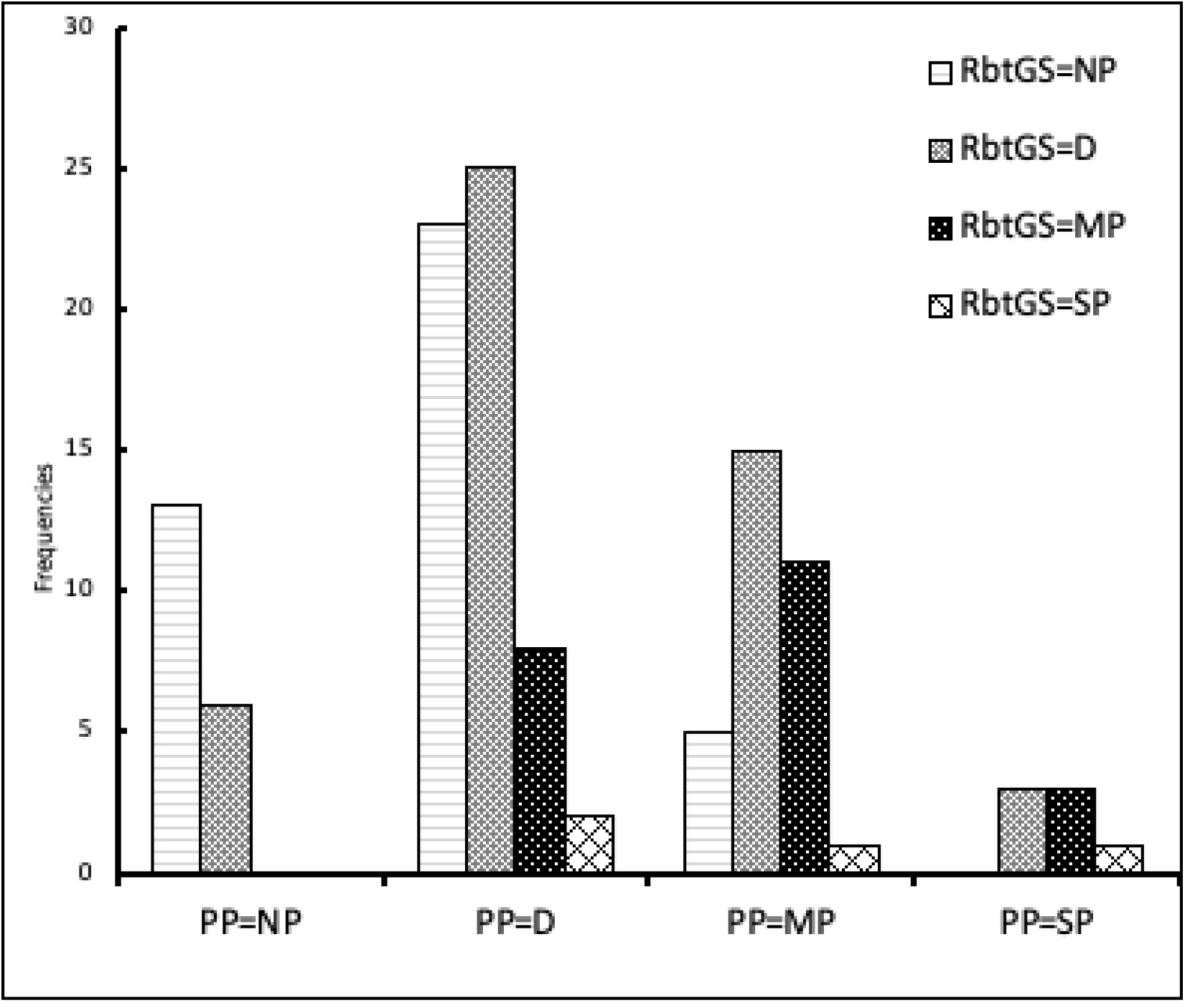
Distribution of the scores resulting from the use of the RbtGS in presumptive pain classes (PP) for group B. Patients in a presumptive condition of pain absence (PP=NP) are mostly classified as NP or D; rabbits in discomfort (PP=D) are mostly classified as NP or D; presumed MP cases are classified as D and MP most of the times; the RbtGS was the only tool that was able to detect some cases of presumed several pain. X^2^ test results (p<0.005) show that frequencies are not randomly obtained, but diagnosis obtained by assessing pain with the RbtGS are related to PP.

## Discussion

Pain assessment and quantification are considered challenging in Veterinary Medicine, due to differences in the expression of pain among species. Moreover, rabbits as a prey species are naturally prone to mask any sign of pain, making the recognition even more difficult^13^. Despite this, pain assessment is essential in order to meet the ethical and clinical necessity to provide analgesia in suffering patients^25–26^. In rabbits there is evidence that pain causes a decrease in activity and appetite^27–28^; since rabbits’ metabolism is geared to a constant supply of nutrients from the digestive tract, a decreased or absent food intake and the subsequent mobilization of fat reserves can lead to ketoacidosis and hepatic lipidosis^5^.

A very recent review article about rabbit analgesia^3^ describes how, according to some studies, veterinarians who doubt their knowledge in pain assessment in cats and dogs are respectively 30% and 42%^29–30^; this percentage raises up to 60% for small mammals such as rabbits and guinea pigs, according to a study that was conducted in New Zealand^31^, and the lack of a ‘gold standard’ method for pain assessment leads to difficulties in pain management. To verify properly the effectiveness of analgesic drugs, a validated assessment tool is needed. Moreover, the drugs used to treat pain in rabbits, were tested on small samples and few papers were published^6,3^. For all these reasons, the effect and the duration of analgesia are often different from what the clinician would expect.

Physiological parameters can be altered in several circumstances such as stress, positive excitement and any pathological condition^13^. Handling during the clinical examination is stressful for the patient, therefore physiological parameters such as respiratory and heart rates have only a limited function in pain assessment^4–5^. Behavioral changes are considered to be a more reliable sign of pain and the return to a calm and unstressed behavior often coincides with pain resolution^32^.

To properly assess the presence of an abnormal behavior, familiarity with the typical behavior of the species, knowledge of the individual characteristics and a prolonged observational period are fundamental. Therefore, it is hard to assess behavioral changes during a relatively short clinical examination.

Pain scales are structured evaluation tools and allow a fast quantification of pain intensity, helping the clinician to determine if pain is present and if there is any variation due to the analgesic treatment^12^. The RbtGS is the only existing validated scale for acute pain evaluation in laboratory rabbits; it is based on five facial action units^13^ that can be observed from a distance in a relatively short time, making the evaluation less stressful for the patient. Although, there are some limitations: the RbtGS is based on one rabbit breed only, NZW, while the pet rabbit population presents several breeds, with many morphological variations; additionally, RbtGS has never been tested in a clinical setting yet, since laboratory rabbits were supposed to be healthy and undergoing a routinely procedure such as ear tattooing; in fact, research findings rarely take into consideration the differences in age and potential concurrent health problems that are more likely to be found in pet rabbits.

The latter considerations underline the necessity to develop a reliable tool to quantify pain, usable in a clinical environment on several breeds, by operators with difference degree of experience. Furthermore, the scale should be fast to use, in order to avoid any delay in the analgesic treatment and it should not be invasive, in order to prevent any further stress for the patient.

This study was intended to develop a multidimensional composite pain scale for pain evaluation in rabbits (CANCRS) and to assess its validity and the validity of the RbtGS in clinical environment on various breeds. Furthermore, our aim was to develop a preliminary pain classes definition, in order to take a further step towards the use of pain scales as a tool do determine whether analgesia is needed. This study has been performed in a clinical setting and serum corticosterone concentrations measurements were not performed, on the contrary on what has been done from Keating et al. (2012)^13^. Results were compared to a presumptive pain diagnosis, according to what is reported literature for other domestic mammals^22^. Although it should be remembered that pain is a subjective experience that is not possible to completely describe except for self-reported pain, possible only in human medicine.

As previously mentioned, composite scales should be reliable and easy to use; for this reason, an assessment of inter-rater reliability between V and S was performed for each parameter using Cohen’s kappa and for each scale using ICC. Inter-observer variation results show that the agreement between V and S raters for all parameters included in the CANCRS are good or very good. Specifically, the FAU included in the RbtGS result to have a good inter-rater agreement, except for ear position, for which Cohen’s kappa resulted to be very good. Although, we cannot exclude that this result is due to the small subject population considered. Independently from their experience degree, the raters resulted equally able to assess all the CPS parameters.

ICC resulted to be very good for both the CANCRS and the CPS, indicating that experience has only a slight influence on the result of the assessment; this is also indicated by the restricted CIs. For what concerns RbtGS, the good ICC and the wider CI, show that rater’s experience has a greater, but still acceptable, impact on the results. Furthermore, our results show that assessing pain in rabbits by using clinical parameters only is not effective; on the contrary, results for CANCRS and RbtGS show that the two scales are efficient in revealing pain conditions. However, pain classes should be revised, since both CANCRS and RbtGS tend to classify as absence of pain (NP) or discomfort (D) even the cases of more intense pain, that should be classified as moderate (MP) or severe pain (SP). Therefore, both CANCRS and RbtGS tend to underestimate pain, according to presumptive pain diagnosis; although, best performances were obtained with the RbtGS, that was able to reveal some cases of severe pain.

It is worth mentioning that patients were evaluated during the clinical examination, but no follow up was performed. The responsiveness of the scale to changes in pain level after analgesic administration was not tested and this represents an area for further study.

Using the CANCRS requires about 3 minutes, which is consistent with a clinical setting use. Since RbtGS is included in the CANCRS, using this alone is faster, but the clinical parameters that the latter scale add to the RbtGS, should always be monitored during any clinical examination; therefore, using the CANCRS can lead to a more exhaustive evaluation of the patient.

Even if clinical parameters resulted to be easy to assess, the scores obtained using the CPS are not related to the presumptive pain diagnosis; this remarks that clinical parameters alone are neither effective or reliable in quantifying pain. The suggestion for a clinician wanting to assess and quantify pain is to use clinical parameters as a support to pain diagnosis otherwise formulated. The RbtGS resulted to be useful in clinical practice, when a wide variety of rabbit breeds is considered; however, reliability results show that training is important for a better use of the scale. The CANCRS was easy and fast to use, showing good inter-rater reliability when used by veterinarians and veterinary medicine students. A refinement of the scale pain classes could be beneficial to increase validity of results, in order to be used as a tool to support the clinician with choosing whether an analgesic plan is needed.

## Acknowledgments

The authors are extremely grateful to Dr Matt Leach from the School of Agriculture, Food & Rural Development, Newcastle University, who provided us with the material and information about the proper use of the RbtGS.

Data Availability Statement: All relevant data are within the paper (and its Supporting information file).

## Supporting information

S1 Dataset.

